# Amelioration of symptomatic Alzheimer’s Disease after selective impairment of p75^NTR^ function in adult forebrain excitatory neurons

**DOI:** 10.1101/2025.10.08.681148

**Authors:** Xuetong Li, Meng Xie, Carlos F. Ibáñez

## Abstract

The p75 neurotrophin receptor (p75^NTR^) contributes to the development of Alzheimer’s Disease (AD) pathology by enhancing amyloid precursor protein (APP) cleavage and amyloid plaque formation. However, the cell type-specific and temporal roles of p75^NTR^ in AD progression remain unclear. Here, we report that conditional knock-in of functionally impaired p75^NTR^ variants lacking the death domain (ΔDD) or transmembrane Cys^259^ (C259A) specifically in forebrain excitatory neurons of 5xFAD mice significantly attenuated multiple AD-associated pathologies, including amyloid plaque accumulation, gliosis, neurite dystrophy, as well as learning and memory deficits. Hippocampal amyloid plaque burden was reduced to levels comparable to those found in whole-body knock-in mice. Strikingly, delaying introduction of p75^NTR^ variants until advanced disease stages produced comparable beneficial effects, and rescued behavior performance in cognitively impaired animals. These findings suggest that blunting p75^NTR^ function can have beneficial effects even during symptomatic stages of AD, offering a potential therapeutic approach complementary to passive vaccination.

## Introduction

Alzheimer’s disease (AD) is the most prevalent form of dementia. The generation and accumulation of neurotoxic fragments of amyloid beta (Aβ), generated from amyloid precursor protein (APP) through sequential proteolytic cleavage, is one primary cause of AD pathogenesis (also known as the amyloid hypothesis) (Beyreuther & Masters, 1991; Selkoe, 1991; Hardy & Allsop, 1991). Forebrain excitatory neurons express significant levels of APP which can be cleaved by the β-site secretase (BACE1) within its extracellular domain (Vassar *et al*, 1999). This cleavage generates a C-terminal fragment (CTFβ) which serves as a substrate for a γ-secretase to produce Aβ peptides of 40 or 42 amino acids, the main component of the AD amyloid plaques (Bitan *et al*, 2003; SELKOE, 2000; Vassar *et al*, 2009). APP can also be cleaved by extracellular α-secretases (e.g. ADAM10) within the Aβ sequence to generate soluble APP α (sAPPα), thereby precluding cleavage by BACE1 and Aβ production. β-site and α-site secretases represent antagonistic processing alternatives of amyloidogenic and non-amyloidogenic cleavage of APP, respectively (Furukawa *et al*, 1996; Meziane *et al*, 1998; Stein *et al*, 2004).

p75 neurotrophin receptor (p75^NTR^), also known as tumor necrosis factor receptor superfamily 16 (TNFRSF16), is a member of the death receptor superfamily characterized by the presence of a 6 α-helix bundle known as the ”death domain” (DD) in its intracellular region (Liepinsh *et al*, 1997). The intracellular domain of p75^NTR^ lacks catalytic activity and participates in signal transduction through ligand-induced recruitment and release of effector proteins (Bronfman & Fainzilber, 2004; Dechant & Barde, 2002; Roux & Barker, 2002; Ibáñez & Simi, 2012). At the cell surface, p75^NTR^ forms dimers stabilized by a disulfide-link formed by a highly conserved cysteine residue (Cys^259^) in its transmembrane domain (Vilar *et al*, 2009). This allows for a ligand-induced conformational change to be propagated from extracellular to intracellular domains that results in the separation of the DDs thereby facilitating the binding of effector proteins that regulate different downstream signaling pathways (Vilar *et al*, 2009). p75^NTR^ can bind all four types of mature neurotrophins as well as their pro-neurotrophin precursors, and has been shown to participate in a wide range of biological processes across diverse cell types, including neuronal death and survival, axonal degeneration and synaptic dysfunction (for reviews see (Underwood & Coulson, 2008; Ibáñez & Simi, 2012; Chao, 2003).

Expression of p75^NTR^ is upregulated upon brain injury and neuronal damage, and many studies have linked p75^NTR^ to neurodegenerative diseases (Ibáñez & Simi, 2012) . Analyses of brains of AD patients and various animal models demonstrate increased p75^NTR^ expression during disease progression (Chakravarthy *et al*, 2012; Hu *et al*, 2002; Mufson & Kordower, 1992). Aβ can directly interact with p75^NTR^ and induce rapid cell death in cultured neurons through activation of downstream apoptotic signaling (Knowles *et al*, 2009; Perini *et al*, 2002; Rabizadeh *et al*, 1994; Sotthibundhu *et al*, 2008; Yaar *et al*, 1997; Tanaka *et al*, 2016), although it remains unclear how this contributes to neuronal death in human AD which can take decades to occur. Null mutation of the gene encoding p75^NTR^ in the 5xFAD mouse model (expressing human APP and PSEN1 transgenes carrying three and two AD-linked mutations, respectively (Oakley *et al*, 2006)) partially alleviates AD pathology (Yi *et al*, 2021). Notably, our laboratory has previously shown that two signaling-deficient p75^NTR^ variants, ΔDD (lacking the DD) and C259A (with Ala replacing the highly conserved transmembrane Cys^259^ in the transmembrane domain), resulted in greater neuroprotective effects than the full knockout of the receptor when introduced in the germline of 5xFAD mice (Yi *et al*, 2021). Mechanistically, this was achieved by the ability of these variants to slow down APP internalization and thereby interfere with the amyloidogenic pathway of APP processing (Yi *et al*, 2021). Because these mutations were introduced directly in the mouse germline, it remained unclear which p75^NTR^-expressing cells (i.e. specific subpopulations of neurons or glia) were responsible for the neuroprotective effects, nor whether any neuroprotection could be achieved if the variants were to be expressed after the overt establishment of AD pathology (i.e. during symptomatic disease stages).

In this study, we conditionally introduced the ΔDD and C259A variants of p75^NTR^ specifically into forebrain excitatory neurons of 5xFAD mice by first generating” floxed” p75^ΔDD^ and p75^C259A^ alleles and subsequently breeding in the CamK2a-Cre driver or its tamoxifen-inducible variant CamK2a-Cre^ERT2^ (Tsien *et al*, 1996). This approach allowed to assess what level of neuroprotection could be achieved through the expression of these variants in a very restricted subset of neurons and whether any neuroprotection could be obtained by substantially delaying the introduction of p75^NTR^ variants to advanced, overtly symptomatic stages of the disease. The results greatly surpassed our expectations, firmly establishing p75^NTR^ as a worthy therapeutic target in AD.

## Results

### Generation of conditional knock-in mice expressing signaling-deficient p75^NTR^ variants in forebrain excitatory neurons

To investigate the effects of signaling-deficient p75^NTR^ variants in a cell type-specific and time-dependent manner, we generated conditional alleles of the mouse *Ngfr* gene encoding p75^NTR^ (herein termed *ΔDD^fl^* and *C259A^fl^*) that allow expression of mutant ΔDD and C259A variants after Cre-mediated recombination (Fig. EV1A, B). We confirmed that the engineered alleles expressed normal levels of p75^NTR^ prior to recombination (Fig. EV2A, B). In order to specifically introduce ΔDD and C259A p75^NTR^ variants into forebrain neurons, we employed the *Camk2a-Cre* driver which induces robust and specific recombination in forebrain excitatory neurons (Tsien *et al*, 1996) . We verified the specificity of this driver by breeding it to Ai14 reporter mice and assessed the proportion of tdTomato^+^ cells among the NeuN^+^ subpopulation. tdTomato expression began at 2 weeks of age, reaching 90% by 2 months in hippocampus and cortex (Fig. EV3A, B), and was restricted to forebrain neurons, as no tdTomato signal was detected in the midbrain or cerebellum, and no overlap was observed between tdTomato and either GFAP or Iba1 in the cortex or hippocampus (Fig. EV3C-E). We generated mice carrying the *Camk2a-Cre* driver and *ΔDD^fl/fl^* and *C259A^fl/fl^*alleles (herein referred to as *ΔDD^Camk2a^* and *C259A^Camk2a^*, respectively). Cre-mediated recombination was verified by PCR from forebrain tissue of 2-month-old mice (Fig. EV4A, B). At the protein level, deletion of the death domain in *ΔDD^Camk2a^* mice was verified by Western blotting of hippocampus extracts using antibodies directed to either the extracellular (ECD) or death domains (DD) (Fig. EV5A, B). Similar results were obtained by immunohistochemistry in cortex and hippocampus of 2 month old *ΔDD^Camk2a^*mice (Fig. EV5C). Together, these results confirm the successful generation of conditional knock-in mice that specifically express signaling-deficient p75^NTR^ variants in forebrain neurons.

### Signaling-deficient p75^NTR^ variants in forebrain excitatory neurons of 5xFAD mice reduce Aβ plaque burden and enhance non-amyloidogenic APP processing

We generated 5xFAD mice expressing signaling-deficient p75^NTR^ variants in forebrain excitatory neurons (termed *ΔDD^Camk2a^*/5xFAD and *C259A^Camk2a^*/5xFAD, respectively) and assessed their amyloid plaque burden in hippocampus and cortex at 6.5 and 9 months by Aβ immunohistochemistry in comparison to control 5xFAD mice carrying non-recombined p75^NTR^ alleles (*ΔDD^fl/fl^*/5xFAD and *C259A^fl/fl^*/5xFAD, respectively). Both *ΔDD^Camk2a^*/5xFAD and *C259A^Camk2a^*/5xFAD mice showed a significant reduction in Aβ plaque burden to approximately 50% of control 5xFAD levels in hippocampus and cortex at both ages and in both male (Fig. 1A-C) and female (Fig. EV6A-C) mice. Importantly, Aβ plaque burden in thalamus remained unchanged (Fig. 1D), in agreement with the specificity of the *Camk2a-Cre* driver. We also noted that the proportion of large Aβ plaques (>50 μm diameter) decreased, while that of small plaques (<30 μm diameter) increased in the brains of *ΔDD^Camk2a^*/5xFAD and *C259A^Camk2a^*/5xFAD mice (Fig. 1E), indicating that Aβ plaque burden was reduced in both number and size by the conditional mutations. Interestingly, Aβ plaque levels in *C259A^Camk2a^*/5xFAD mice was comparable to those observed in constitutive, whole-body p75^C259A^/5xFAD mice (Fig. 1A, B), suggesting that restricted expression of this p75^NTR^ variant in only postnatal forebrain excitatory neurons can confer similar neuroprotection to that afforded by whole-body expression from birth (as that reported in our earlier study by Yi et al 2021).

**Fig. 1.**
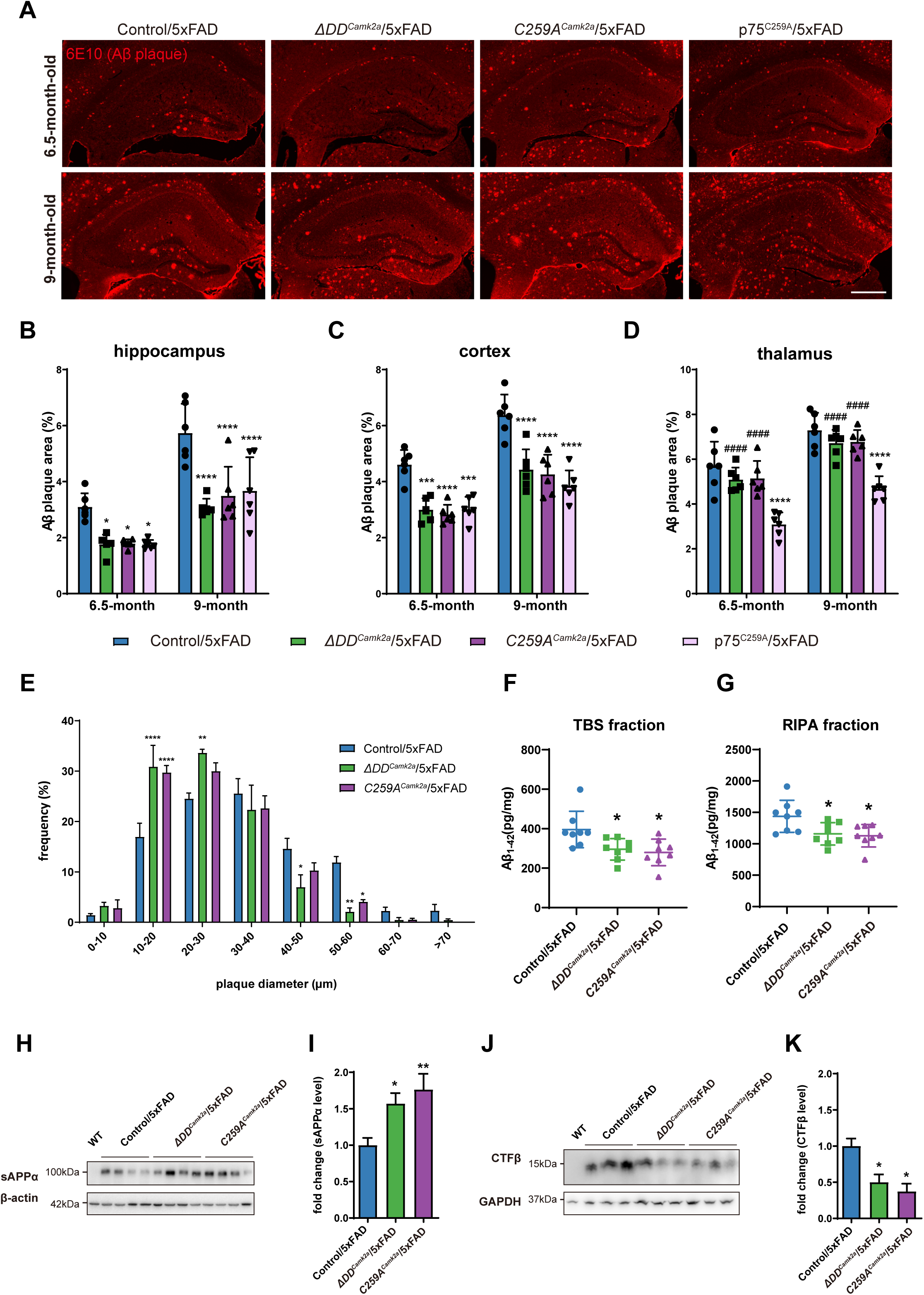
**Signaling-deficient p75^NTR^ variants in forebrain excitatory neurons of 5xFAD mice reduce Aβ plaque burden and enhance non-amyloidogenic APP processing** (A) Immunostaining of Aβ plaques with 6E10 antibody in the hippocampus of Control/5xFAD, ΔDD^Camk2a^/5xFAD, C259A^Camk2a^/5xFAD and p75^C259A^/5xFAD mice of the indicated ages. Scale bar, 500 μm. (B-D) Quantification of Aβ plaque burden in the hippocampus (B), cortex (C) and thalamus (D) of Control/5xFAD, ΔDD^Camk2a^/5xFAD, C259A^Camk2a^/5xFAD and p75^C259A^/5xFAD mice as indicated. Histogram shows the percentage of hippocampal area occupied by Aβ plaques (mean±SD, N=6 animals per group). All mice included in this analysis were male. Two-way ANOVA followed by Tukey’s multiple comparisons test. *p<0.05, **p<0.01, ***p<0.001, ****p<0.0001 versus Control/5xFAD. ####, p<0.0001 versus p75^C259A^/5xFAD. (E) Frequency distribution analysis for the plaque diameter in the hippocampus of 6.5-month-old Control/5xFAD, ΔDD^Camk2a^/5xFAD and C259A^Camk2a^/5xFAD mice (mean±SEM, N=3 animals per group). For each animal, more than 300 plaques in the hippocampus are quantified. Two-way ANOVA followed by Tukey’s multiple comparisons test. (F, G) Quantitative ELISA determinations of Aβ1-42 in hippocampus of 9-month-old Control/5xFAD, ΔDD^Camk2a^/5xFAD and C259A^Camk2a^/5xFAD mice. TBS fraction (F) refers to the soluble fraction after Tris-buffered saline extraction and contains Aβ monomers. RIPA fraction (G) refers to the soluble fraction after RIPA buffer extraction from the Tris-buffered Saline pellet and contains oligomeric Aβ. Shown is mean±SEM, N=8 animals per group. One-way ANOVA followed by Tukey’s multiple comparisons test. All mice included in this analysis were male. (H) Western blot analysis of soluble APP alpha (sAPPα) in the Tris-buffered saline fraction of hippocampal extraction from 9-month-old 5xFAD mice carrying conditional knock-in p75^NTR^ variants. sAPPα was detected by 6E10 antibody. The lower panel shows β-actin as a control for equal loading amount. (I) Quantification of sAPPα (mean±SD) normalized to internal control and expressed relative to levels in Control/5xFAD. N=8 animals per group. (J) Western blot analysis of CTF beta (CTFβ) in the RIPA fraction of hippocampal extraction from 9-month-old 5xFAD mice carrying conditional knock-in p75^NTR^ variants. CTFβ was detected by 6E10 antibody. The lower panel shows GAPDH as a control for equal loading amount. (K) Quantification of CTFβ (mean±SD) normalized to internal control and expressed relative to levels in Control/5xFAD. N=3 animals per group.

Next, we quantified Aβ1-42 levels by ELISA after differential detergent extraction from hippocampus of 9-month-old mice and found that both *ΔDD^Camk2a^*/5xFAD and *C259A^Camk2a^*/5xFAD mice exhibited lower levels of Aβ1-42 monomers (assessed in the supernatant of the TBS fraction) and oligomers (supernatant of the RIPA fraction) compared to their respective control 5xFAD mice (Fig. 1F, G). Finally, we assessed the levels of APP fragments soluble APP alpha (sAPPα) and carboy-terminal fragment beta (CTFβ) in hippocampal lysates from 9-month-old mice and found increased sAPPα levels (Fig. 1H, I) but decreased CTFβ levels (Fig. 1J, K) in *ΔDD^Camk2a^*/5xFAD and *C259A^Camk2a^*/5xFAD mice compared to 5xFAD controls, indicating a bias towards non-amyloidogenic APP processing in the mutant mice. Together, these results demonstrate that interfering with p75^NTR^ function only in forebrain excitatory neurons is sufficient to reduce Aβ plaque deposition to levels comparable to a systemic intervention.

### Reduced brain AD histopathology in 5xFAD mice expressing signaling-deficient p75^NTR^ variants in forebrain neurons

Accumulation of Aβ plaques induces the activation of astrocytes and microglia which can lead to neuroinflammation and synaptic loss (Deng *et al*, 2024; Strooper & Karran, 2016; Singh, 2022). As expected, astrogliosis and microgliosis were significantly increased at 6 and 9 months of age in hippocampus of control 5xFAD mice (*ΔDD^fl/fl^*/5xFAD and *C259A^fl/fl^*/5xFAD), as shown by increased immunostaining for glial fibrillary acidic protein (GFAP) and Ionized calcium-binding adaptor molecule 1 (Iba1) compared to non-AD mice (Fig. 2A, B). Activated astrocytes and microglia also showed morphological changes and clustered around Aβ plaques. In contrast, 5xFAD mice expressing signaling-deficient p75^NTR^ variants in forebrain neurons showed significantly reduced astrocyte and microglia activation (Fig. 2A-D). Vimentin-expressing astrocytes are a subset of pro-inflammatory, disease-associated cells that contribute to amplify Aβ toxicity promoting neurodegeneration (Habib *et al*, 2020; Lee & Quintana, 2024). Vimentin-expressing astrocytes were detected only in 5xFAD brains and were concentrated in the dentate gyrus area and along the hippocampal border (Fig. 2E). The hippocampus of *ΔDD^Camk2a^*/5xFAD and *C259A^Camk2a^*/5xFAD mice showed approximately 50% reduction in vimentin expression compared to control 5xFAD mice (Fig. 2E, F), suggesting reduced neuroinflammation after disruption of p75^NTR^ signaling in forebrain neurons. Finally, we assessed the accumulation of reticulon 3 (RTN3), a protein that marks degenerating neurites (Hu *et al*, 2007), in cortex and hippocampus of 5xFAD mice expressing different p75^NTR^ alleles. At 9 months of age, control 5xFAD mice exhibited pronounced RTN3 immunostaining clustering around Aβ plaques in both cortex and hippocampus (Fig. 2G). In contrast, RTN3 immunoreactivity was significantly reduced in both *ΔDD^Camk2a^*/5xFAD and *C259A^Camk2a^*/5xFAD mice (Fig. 2G, H). Together, these data indicate reduced gliosis and neurodegeneration in the forebrain of 5xFAD mice expressing signling-deficient p75^NTR^ variants, in agreement with their decreased Aβ burden.

**Fig. 2.**
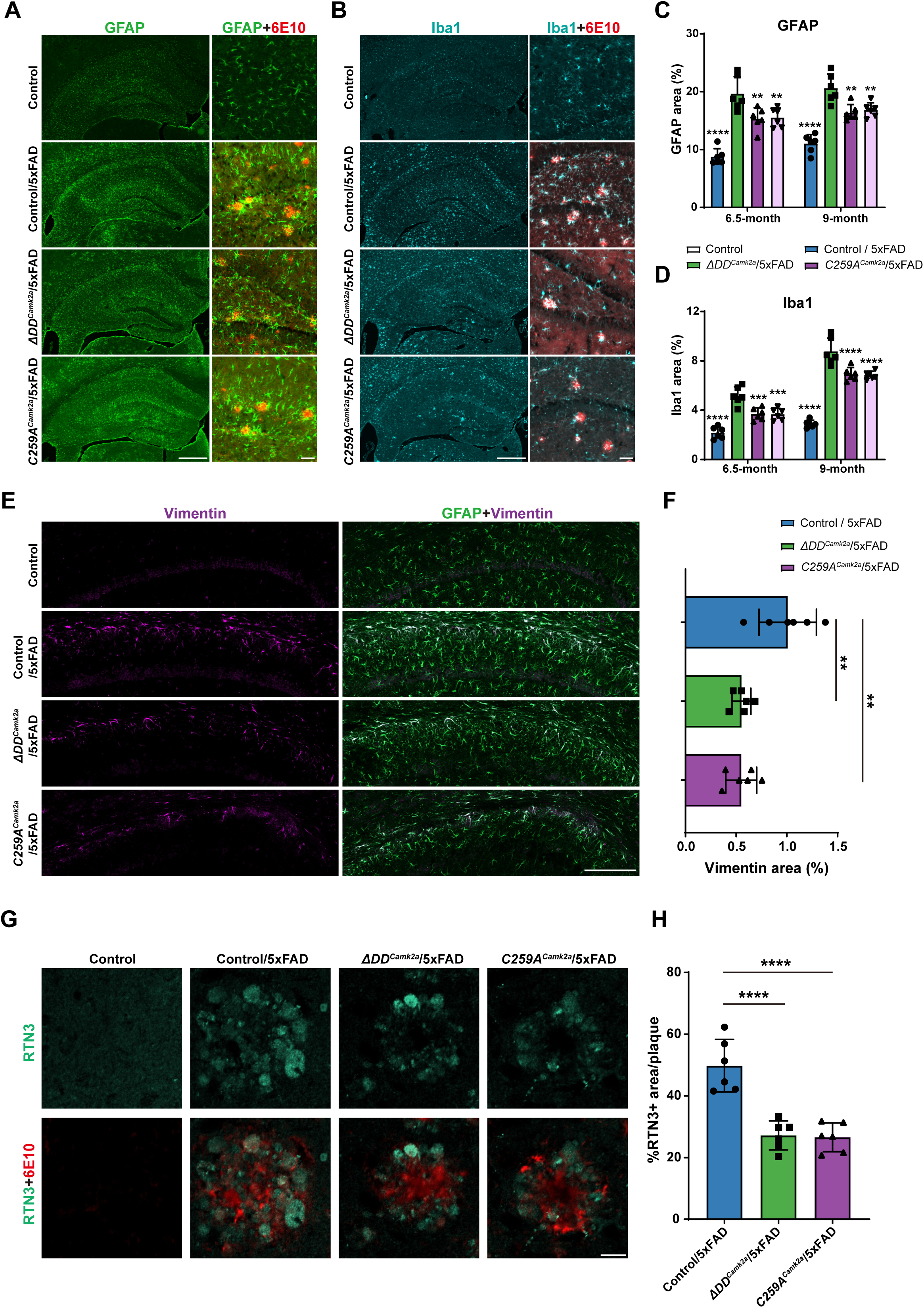
**Reduced brain AD histopathology in 5xFAD mice expressing signaling-deficient p75^NTR^ variants in forebrain neurons** (A) Immunostaining of glial fibrillary acidic protein (GFAP) in the hippocampus of 6.5-month-old Control, Control/5xFAD, ΔDD^Camk2a^/5xFAD and C259A^Camk2a^/5xFAD mice. Scale bar, 500 μm. Right panels show co-immunostaining of GFAP and Aβ plaques at a higher magnification. Scale bar, 50 μm. (B) Immunostaining of Ionized calcium binding adaptor molecule 1 (Iba1) in the hippocampus of 6.5-month-old Control, Control/5xFAD, ΔDD^Camk2a^/5xFAD and C259A^Camk2a^/5xFAD mice. Scale bar, 500 μm. Right panels show co-immunostaining of Iba1 and Aβ plaques at a higher magnification. Scale bar, 50 μm. (C) Quantification of astrogliosis reflected by GFAP immunostaining in the hippocampus of different mouse strains as indicated. Histograms shows the percentage of hippocampal area occupied by GFAP immunostaining (mean±SD, N=6 mice per group). Two-way ANOVA followed by Tukey’s multiple comparisons test. **p<0.01, ****p<0.0001 versus Control/5xFAD. All mice included in this analysis were male. (D) Quantification of microgliosis reflected by Iba1 immunostaining in the hippocampus of different mouse strains as indicated. Histograms shows the percentage of hippocampal area occupied by Iba1 immunostaining (mean±SD, N=6 mice per group). Two-way ANOVA followed by Tukey’s multiple comparisons test. ***p<0.001, ****p<0.0001 versus Control/5xFAD. All mice included in this analysis were male. (E) Immunostaining of GFAP and Vimentin in the hippocampus of 9-month-old Control, Control/5xFAD, ΔDD^Camk2a^/5xFAD and C259A^Camk2a^/5xFAD mice. Scale bar, 500 μm. (F) Quantification of Vimentin immunostaining in the hippocampus of different mouse strains as indicated. Histograms shows the percentage of hippocampal area occupied by Vimentin immunostaining (mean±SD, N=6 mice per group). One-way ANOVA followed by Tukey’s multiple comparisons test. ***p<0.001, ****p<0.0001 versus Control/5xFAD. All mice included in this analysis were male. (G) Immunostaining of reticulon 3 (RTN3) and Aβ plaques in the hippocampus of 9-month-old Control, Control/5xFAD, ΔDD^Camk2a^/5xFAD and C259A^Camk2a^/5xFAD mice. Scale bar, 15 μm. (H) Quantification of RTN3-positive dystrophic neurites in the hippocampus of different mouse strains as indicated. Histograms shows the percentage of Aβ plaque area that overlapped with RTN3 immunostaining (mean±SD, N=6 mice per group). One-way ANOVA followed by Tukey’s multiple comparisons test. ***p<0.001, ****p<0.0001 versus Control/5xFAD. All mice included in this analysis were male.

### Improved learning and memory in 5xFAD mice expressing ΔDD and C259A p75^NTR^ variants in forebrain neurons

5xFAD mice show significant learning and memory impairments at 6 months of age (Forner et al., 2021). We evaluated learning and memory in 6 month old 5xFAD male mice expressing different p75^NTR^ alleles using the Barnes maze, a widely used test of hippocampal-dependent spatial learning (Bach *et al*, 1995; Barnes, 1979), and the Novel Object Recognition (NOR) test, which assesses the mouse’s natural preference to explore a novel object over a familiar one. During the training phase of the Barnes maze, wild type mice demonstrated normal learning ability, gradually reducing their latency to enter the escape tunnel to about 40s (Fig. 3A). In contrast, control 5xFAD mice (both *ΔDD^fl/fl^*/5xFAD and *C259A^fl/fl^*/5xFAD) showed impaired learning, with essentially constant escape latencies throughout the training sessions (Fig. 3A). Importantly, learning ability was significantly rescued in both *ΔDD^Camk2a^*/5xFAD and *C259A^Camk2a^*/5xFAD mice, with escape latencies reaching wild type levels by the last training session (Fig. 3A). In the memory test conducted 3 hours after the last training session, wild type mice and 5xFAD mice expressing mutant p75^NTR^ variants spent most of their exploration time in the target quadrant containing the escape tunnel, while control 5xFAD mice spent only approximately 25%, consistent with chance probability (Fig. 3B), suggesting improved spatial memory in *ΔDD^fl/fl^*/5xFAD and *C259A^fl/fl^*/5xFA mice. In the training session of the NOR test, no preference for any of the two (identical) objects was observed in any line of mice, as expected (Fig. 3C). In probe tests performed 3 and 24 hours after the training, wild type mice spent approximately 40% more time exploring the new object (Fig. 3D, E). Control 5xFAD mice showed no preference, while both *ΔDD^fl/fl^*/5xFAD and *C259A^fl/fl^*/5xFA mice demonstrated a trend towards recognising the novel object more frequently, although statistical significance was not reached for these groups (Fig. 3D, E).

**Fig. 3.**
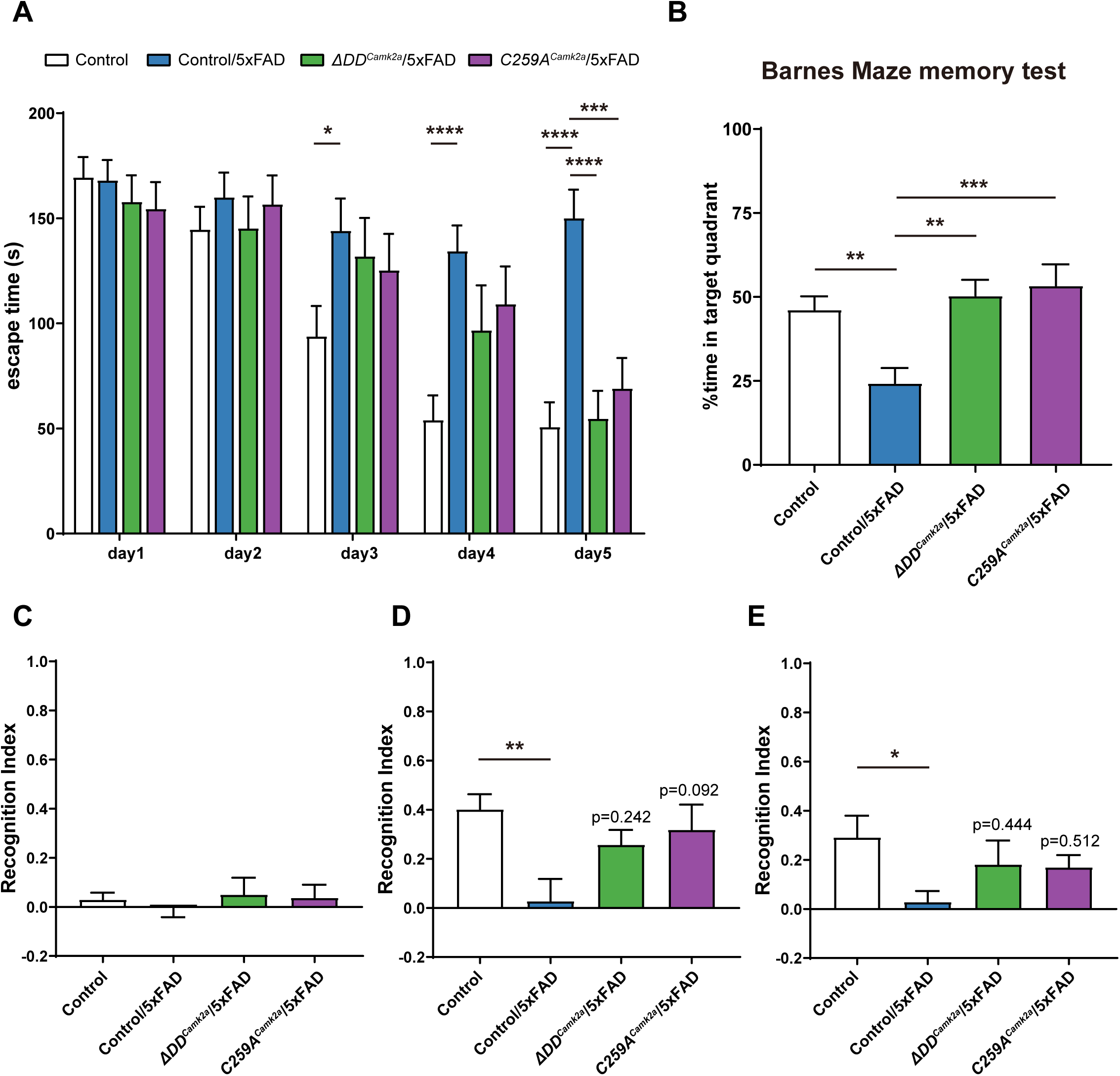
**Improved learning and memory in 5xFAD mice expressing ΔDD and C259A p75^NTR^ variants in forebrain neurons** (A) Training latency in the Barnes maze test of 6-month-old Control, Control/5xFAD, ΔDD^Camk2a^/5xFAD and C259A^Camk2a^/5xFAD mice as indicated. Histograms show the escape time to find the tunnel in the training session of 5 consecutive days (mean±SEM). N=17 (control and Control/5xFAD), 12 (ΔDD^Camk2a^/5xFAD) 14 (C259A^Camk2a^/5xFAD) mice per genotype, respectively. Two-way ANOVA followed by Tukey’s multiple comparisons test. *p<0.05, **p<0.01, ***p<0.001, ****p<0.0001 versus Control/5xFAD. All mice included in this analysis were male. (B) Percentage of time (mean±SEM) spent in the target quadrant of the Barnes maze test 3h after training. One-way ANOVA followed by Tukey’s multiple comparisons test. *p<0.05, **p<0.01 versus Control/5xFAD. (C-E) Behavior performance in the novel object recognition (NOR) test of 6-month-old Control, Control/5xFAD, ΔDD^Camk2a^/5xFAD and C259A^Camk2a^/5xFAD mice as indicated. Histograms show recognition index (mean±SEM) during training (C), 3h after training (D, referred to as short-term memory), and 24h after training (E, referred to as long-term memory). N=12 (control), 15 (Control/5xFAD), 8 (ΔDD^Camk2a^/5xFAD) 8 (C259A^Camk2a^/5xFAD) mice per genotype, respectively. Two-way ANOVA followed by Tukey’s multiple comparisons test. *p<0.05, **p<0.01 versus Control/5xFAD. All mice included in this analysis were male.

### Significant amelioration of AD neuropathology and cognitive function after acute disruption of p75^NTR^ activity during symptomatic AD stages

The observation that impaired p75^NTR^ function in forebrain excitatory neurons provided significant neuroprotection in 5xFAD mice prompted us to investigate whether beneficial effects could still be observed if the mutations were to be introduced after the onset of AD symptoms in these mice. To this end, we used the *Camk2a-CreERT2* driver, which can induce recombination in forebrain neurons after administration of tamoxifen (Madisen *et al*, 2010). We first verified neuron-specific recombination by immunostaining for NeuN, GFAP and Iba1 after tamoxifen administration in 2, 4 and 6 month old *Camk2a-CreERT2*;Ai14 reporter mice (Fig. EV7A-C). Recombination rates were comparable across different injection ages, reaching 70-80% in the hippocampus and 50-70% in the cortex (Fig. EV7D, E). In line with a previous report (Erdmann *et al*, 2007), approximately 20% recombination was observed in neurons of mice injected with vehicle (Fig. EV7D-F).

We introduced the *Camk2a-CreERT2* driver in *ΔDD^fl/fl^*/5xFAD and *C259A^fl/fl^*/5xFAD mice and assessed AD neuropathology in their progeny, herein referred to as *ΔDD^CreERT2^*/5xFAD and *C259A^CreERT2^*/5xFAD mice, respectively. Tamoxifen was administered at 2, 4 or 6 months and analyses were conducted at 9 months of age (Fig. 4A), with the 2 months time point being comparable to the non-inducible *Camk2a-Cre* driver, which achieves maximal Cre recombination at this age (see above). To our surprise, we found comparable reduction in Aβ plaque levels in both hippocampus and cortex at all injection ages (Fig. 4B-F), suggesting that acute disruption of p75^NTR^ activity at advanced AD stages can still afford significant neuroprotective effects. In agreement with this, astrogliosis, as assessed by GFAP immunostaining, was markedly decreased in both *ΔDD^CreERT2^*/5xFAD and *C259A^CreERT2^*/5xFAD mice at all time points of tamoxifen treatment (Fig. 4G, H). Interestingly, microgliosis, as assessed by Iba1 immunostaining, was only reduced in the mice that received tamoxifen at 2 months of age (Fig. 4I, J). To assess whether the effects observed at the tissue level had an impact on cognitive performance, we compared 5xFAD mice expressing p75^NTR^ variants at different times of the disease process in the Barnes maze. Strikingly, both *ΔDD^CreERT2^*/5xFAD and *C259A^CreERT2^*/5xFAD mice showed improved learning ability compared to corn oil-injected controls during training, with no significant differences among the 2, 4 and 6-month injection groups (Fig. 5A, B). In the probe test, all three treatment groups spent significantly more time in the target quadrant, indicating improved spatial memory (Fig. 5C, D). Together, these results demonstrate that acute disruption of p75^NTR^ activity during advanced disease stages, i.e. after Aβ plaque formation and onset of cognitive impairment, is still able to confer beneficial effects to AD progression.

**Fig. 4.**
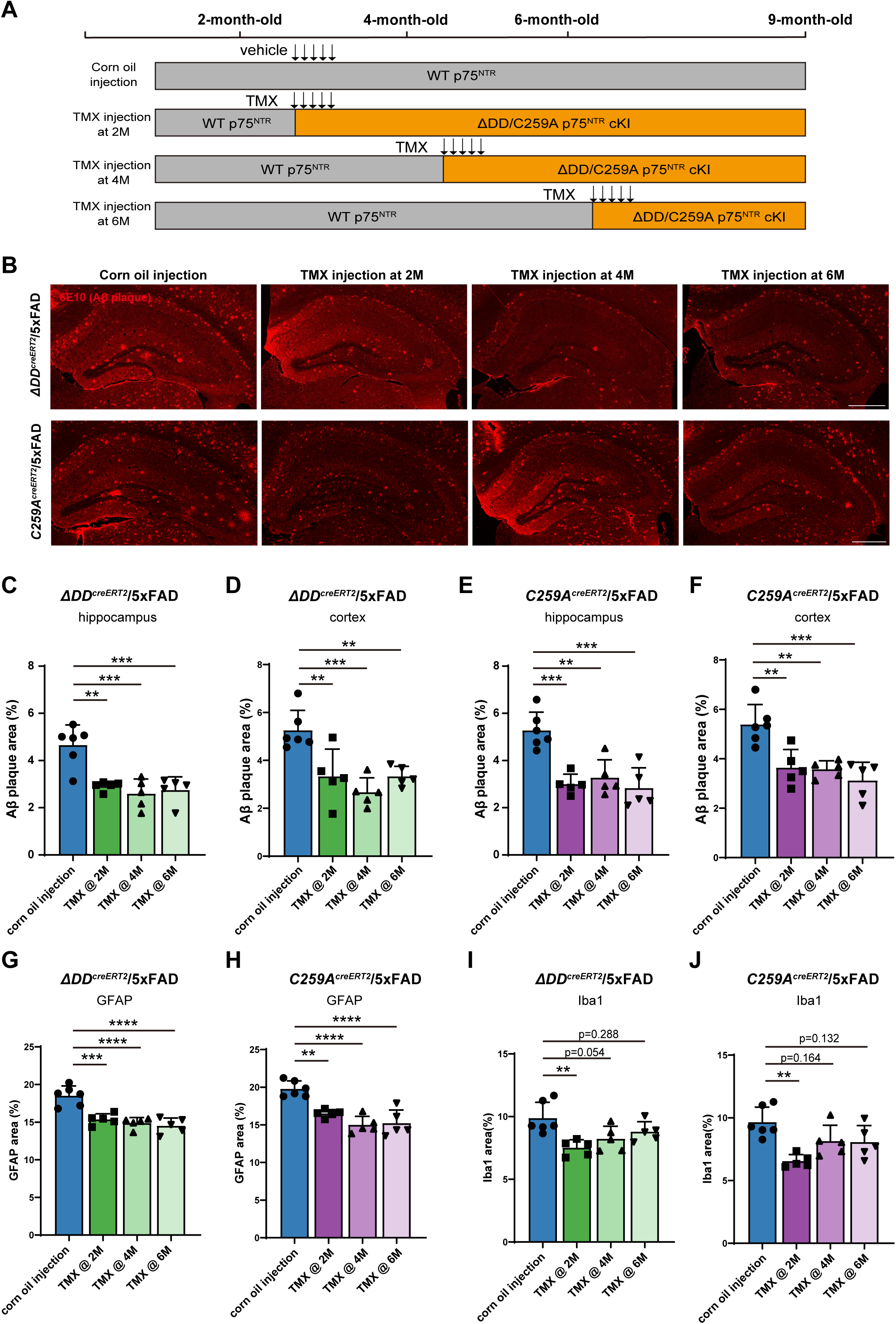
**Significant amelioration of AD neuropathology after acute disruption of p75^NTR^ activity during symptomatic AD stages** (A) Schematic illustration of Tamoxifen administration paradigms. Tamoxifen was administered at 2, 4 or 6 months and analyses were conducted at 9 months of age. (B) Immunostaining of Aβ plaques with 6E10 antibody in the hippocampus of ΔDD^CreERT2^/5xFAD and C259A^CreERT2^/5xFAD 9 month old mice treated with TMX at the indicated ages. Scale bar, 500 μm. (C-D) Quantification of Aβ plaque burden in the hippocampus (C) and cortex (D) of ΔDD^CreERT2^/5xFAD treated with corn oil or TMX as indicated. Histogram shows the percentage of hippocampal or cortical area occupied by Aβ plaques (mean±SD, N=5-6 animals per group). All the mice assessed are males. One-way ANOVA followed by Tukey’s multiple comparisons test. **p<0.01, ***p<0.001 versus Corn oil injected mice. (E-F) Quantification of Aβ plaque burden in the hippocampus (E) and cortex (F) of C259A^CreERT2^/5xFAD treated with corn oil or TMX as indicated. Histogram shows the percentage of hippocampal or cortical area occupied by Aβ plaques (mean±SD, N=5-6 animals per group). All the mice assessed are males. One-way ANOVA followed by Tukey’s multiple comparisons test. **p<0.01, ***p<0.001 versus Corn oil injected mice. (G-H) Quantification of astrogliosis reflected by GFAP immunostaining in the hippocampus of ΔDD^CreERT2^/5xFAD (G) and C259A^CreERT2^/5xFAD (H) treated with corn oil or TMX as indicated. Histograms shows the percentage of hippocampal area occupied by GFAP immunostaining (mean±SD, N=5-6 mice per group). One-way ANOVA followed by Tukey’s multiple comparisons test. **p<0.01, ****p<0.0001 versus Corn oil injected mice. All mice included in this analysis were male. (I-J) Quantification of microgliosis reflected by Iba1 immunostaining in the hippocampus of ΔDD^CreERT2^/5xFAD (I) and C259A^CreERT2^/5xFAD (J) treated with corn oil or TMX as indicated. Histograms shows the percentage of hippocampal area occupied by Iba1 immunostaining (mean±SD, N=6 mice per group). One-way ANOVA followed by Tukey’s multiple comparisons test. ***p<0.001 versus Corn oil injected mice. All mice included in this analysis were male.

**Fig. 5.**
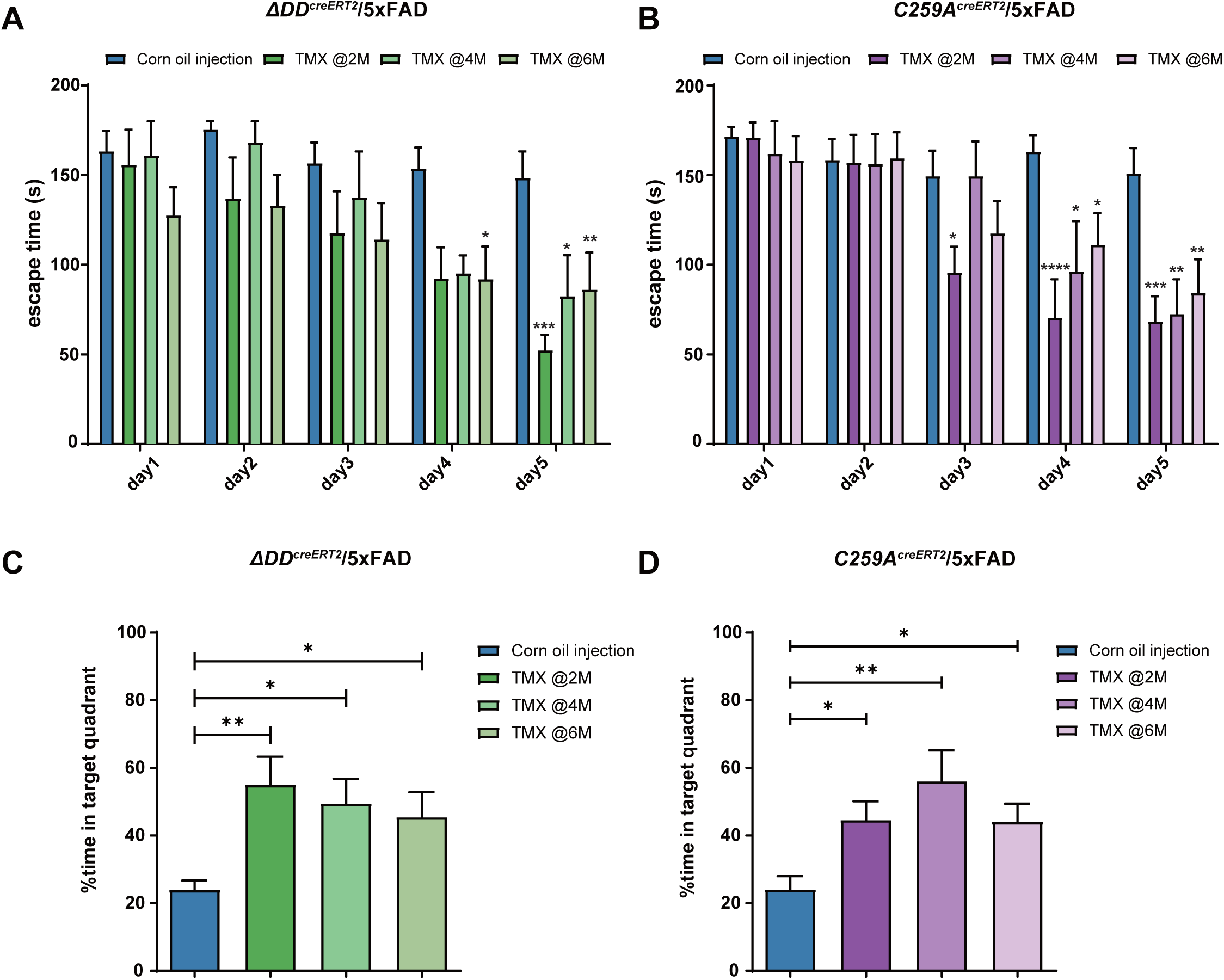
**Significant improvement in cognitive function after acute disruption of p75^NTR^ activity during symptomatic AD stages** (A, B) Training latency in the Barnes maze test of 9-month-old ΔDD^CreERT2^/5xFAD (A) and C259A^CreERT2^/5xFAD (B) treated with corn oil or TMX as indicated. Histograms show the escape time to find the tunnel in the training session of 5 consecutive days (mean±SEM). In panel A, N=13 (corn oil injection), 7 (TMX @2M), 7 (TMX @4M) and 12 (TMX @6M) mice per genotype, respectively. In panel B, N=15 (corn oil injection), 10 (TMX @2M), 6 (TMX @4M) and 10 (TMX @6M) mice per genotype, respectively. Two-way ANOVA followed by Tukey’s multiple comparisons test. *p<0.05, **p<0.01, ***p<0.001, ****p<0.0001 versus Corn oil injected mice. Both male and female mice are used. (C, D) Percentage of time (mean±SEM) spent in the target quadrant 3h after the last training of 9-month-old ΔDD^CreERT2^/5xFAD (C) and C259A^CreERT2^/5xFAD (D) treated with corn oil or TMX as indicated. One-way ANOVA followed by Tukey’s multiple comparisons test. *p<0.05, **p<0.01 versus Corn oil injected mice

## Discussion

Different forms of inactivation of the *Ngfr* gene, from null to point mutations to partial deletion, have been reported to show various degrees of neuroprotective effects in Aβ-based mouse models of AD when introduced constitutively during gestation (Yi *et al*, 2021; Knowles *et al*, 2009; Wang *et al*, 2011). As p75^NTR^ is expressed in several different cell types in the brain, e.g. neurons, glial cells, endothelial cells, it has been unclear whether the beneficial effects afforded by its loss of function arise from all cell types or only one, and if so which one. Moreover, for therapeutic approaches targeting p75^NTR^ to be viable in AD patients, any form of interference with its activity needs to demonstrate beneficial effects during symptomatic stages of the disease, as this is the main cohort of patients that is normally recruited to clinical trials of AD. In this study, we have shown that replacement of native p75^NTR^ with signaling-impaired variants of the receptor in forebrain excitatory neurons is sufficient to significantly alleviate neuropathological and behavioral outcomes in 5xFAD mice, one of the most aggressive mouse models of AD. Although the benefits attained in forebrain structures, namely cerebral cortex and hippocampus, were comparable to those reached by a constitutive mutation from gestation, no reductions in Aβ plaque deposition were observed in the thalamus of either *ΔDD^Camk2a^*/5xFAD or *C259A^Camk2a^*/5xFAD mice, indicating that the effects of these mutations on amyloid burden are brain region-autonomous. Significantly, however, spatial learning and memory (and to some degree also novelty recognition) were rescued to levels similar to those displayed by healthy, non-AD mice, in line with the importance of forebrain structures in these cognitive functions.

Our previous studies showed that signaling-deficient ΔDD and C259A p75^NTR^ variants display significantly reduced internalization and increased partition to recycling endosomes in hippocampal neurons (Yi *et al*, 2021). One consequence of this is that other transmembrane proteins with an ability to interact and co-internalize with p75^NTR^, such as APP, become similarly affected. Thus, both ΔDD and C259A p75^NTR^ delay APP internalization thereby enhancing its availability to plasma membrane α-secretases at the expense of BACE1 β-secretase, which predominantly operates in acidic intracellular compartments (Das *et al*, 2013; Yi *et al*, 2021), leading to reduced Aβ production. In the present study, we found that selective expression of signaling-deficient ΔDD and C259A p75^NTR^ variants in forebrain excitatory neurons significantly reduced the accumulation of the CTFβ fragment, a product of APP cleavage by BACE1, while increasing sAPPα levels, which is generated by α-secretases, in the hippocampus of 5xFAD mice, indicating enhanced non-amyloidogenic APP processing. Thus, mechanistically, the reduction in Aβ levels observed after selective expression of ΔDD and C259A p75^NTR^ variants in forebrain excitatory neurons of 5xFAD mice is likely the result of impaired APP intracellular trafficking, as demonstrated in our previous study (Yi *et al*, 2021).

With regards to the temporal aspects of interrupting normal p75^NTR^ function in AD, a striking result of our study is the significant amelioration of AD neuropathology and cognitive function achieved after acute disruption of p75^NTR^ activity during symptomatic AD stages, particularly at 6 months of age, a stage at which the 5xFAD model show pronounced levels of amyloid deposition, gliosis and behavioral impairment. This finding is highly significant, as all current clinical trials for AD enroll patients that already show symptoms of cognitive impairment, and supports the development of therapeutic strategies targeting p75^NTR^ in AD. Encouragingly, a recent Phase II clinical trial in AD patients of a small molecule presented as a “modulator” of p75^NTR^ activity resulted in a measurable improvement in several AD biomarkers as well as cognitive functions, although the latter did not reach statistical significance (Shanks *et al*, 2024). It is currently unknown how this drug affects p75^NTR^ signaling. In this regard, the results of our present study, particularly the effects of the C259A mutation in the transmembrane domain, provide a new direction for the development of molecules that tweak the activity of p75^NTR^ in a beneficial way, perhaps by targeting the transmembrane domain of the receptor, as shown in previous work from our laboratory (Goh *et al*, 2018; Lopes-Rodrigues & Ibáñez, 2025) .

Unexpectedly, a comparable reduction in Aβ burden was observed in 9 month old *ΔDD^CreERT2^*/5xFAD and *C259A^CreERT2^*/5xFAD mice (roughly 50% of control 5xFAD) regardless of the time of TMX injection (whether 2, 4 or 6 months), suggesting that acute impairment of p75^NTR^ function in symptomatic AD does not reduce amyloid accumulation at a constant rate but up to a certain threshold level, roughly equivalent to that of 6 month old control 5XFAD mice, covering approximately 3% of the hippocampus. Indeed, the level of amyloid accumulated at 9 months in the conditional mutants that received TMX at 6 months was similar to the levels that control 5xFAD mice showed at 6 months, suggesting amyloid levels were stabilized at 6 months after TMX administration, staying constant for another 3 months. Intriguingly, despite comparable levels of amyloid burden, 6 month old 5xFAD mice were cognitively impaired (Fig. 3) but 9 month conditional mutants that received TMX at 6 months were not (or much less so, Fig. 5), suggesting a recovery of cognitive function that cannot solely be explained by Aβ plaque levels in the brain. Although microgliosis remained unchanged in conditional mutants that received TMX at 6 months, astrogliosis was reduced below the level of 6 month old 5xFAD mice, but whether this was a contributing factor to their improved cognitive behavior is unclear at present. Unlike Aβ plaque burden and astrogliosis, only the conditional mutants that received TMX at 2 months exhibited improved microgliosis when examined at 9 months, suggesting that microgliosis requires a longer time to resolve upon reduction of Aβ plaque accumulation.

In conclusion, our demonstration that neuron-specific impairment of p75^NTR^ function confers neuroprotection in a very aggressive model of AD —even at symptomatic stages— positions p75^NTR^ as a promising therapeutic target and opens new avenues for interventions that can be beneficial beyond early disease stages.

## Methods

### Mice

Animal care and experimental procedures were approved by Laboratory Animal Welfare and Ethics Committee of Chinese Institute for Brain Research (CIBR-IACUC-028). Mice were housed in a 12-h light–dark cycle and fed a standard chow diet. The mouse lines utilized in this study are as follows: 5xFAD (Oakley et al, 2006); Camk2a-Cre, JAX Laboratory (#005359); Camk2a-CreERT2, JAX Laboratory, (#012362); conditional knock-in p75^NTR^ ΔDD^fl^ and C259A^fl^ generated for this study by Cyagen Co., China; constitutive knock-in p75^C259A^ (Tanaka *et al*, 2016) . All mice were male of C57BL/6J background.

### Genomic DNA extraction and PCR

The Biotime Genomic DNA Extraction Kit was used to extract genomic DNA from frozen tissue of mice of different genotypes. Electrophoretic results showed bands at 760 bp corresponding to the recombined flox allele; and bands at 2,000 bp corresponding to the unrecombined flox allele. Primers sequences for PCR are as follows:

C259A-flox-F : 5’-GCCTCTGATCAGCTCTAAAAGTGCC-3’

C259A-flox-R : 5’-ATCCTCAGTGTGCCCCCTCTTAC-3’ ΔDD-flox-F: 5’-GGACACTTGCTGCAGTTATAAC-3’ ΔDD-flox-R: 5’-ACAGAGGCCCTACACAGAGAT-3’

### Antibodies

The primary antibodies used for immunofluorescence (IF) and immunoblotting (WB) were as follows:

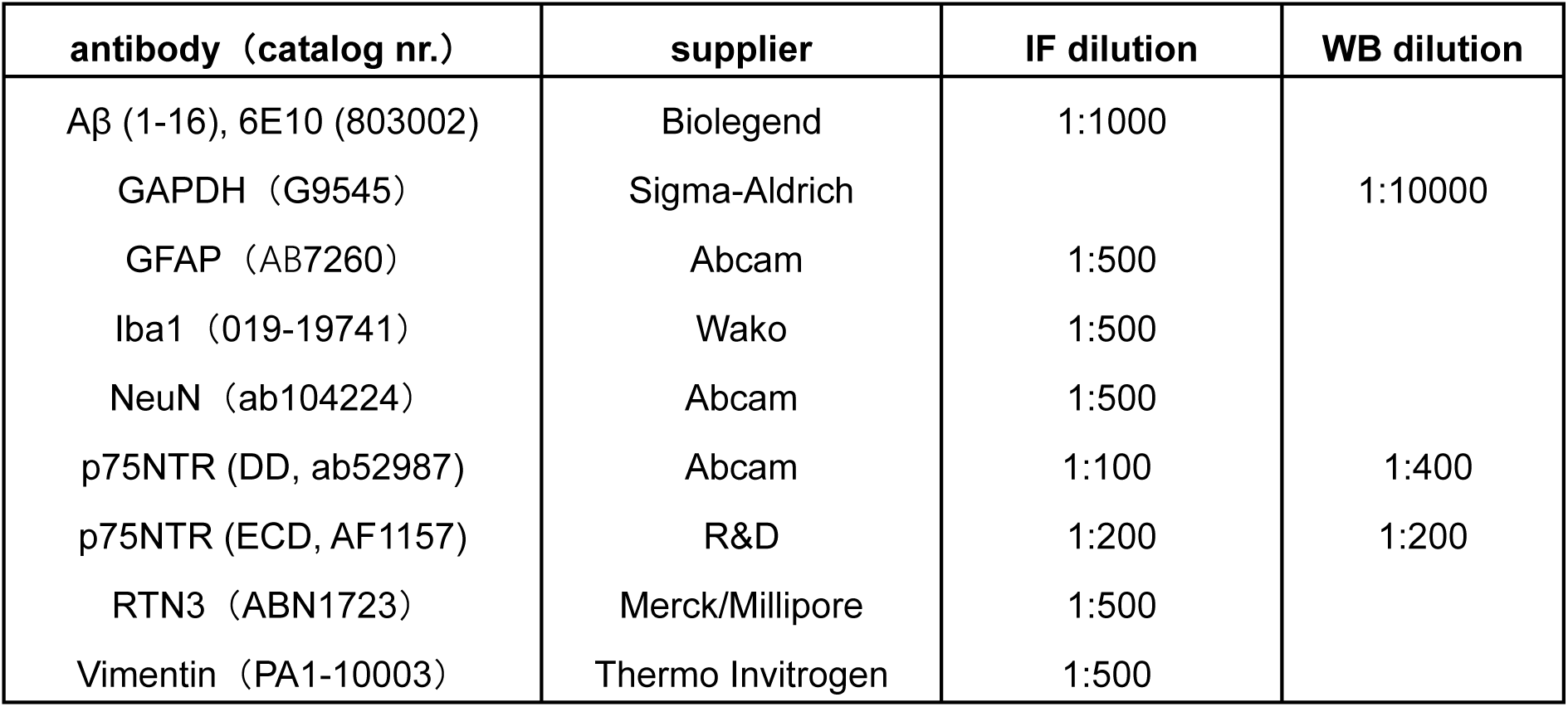

### Western blotting

Frozen tissue was collected and thawed on ice, the tissue was minced and an appropriate amount of RIPA Buffer was added, and the tissue was lysed on ice for 30 min. The lysed samples were centrifuged at 12,000 g for 10 min at 4°C, and the supernatant was taken. The protein concentration contained in the samples was determined using a BCA kit (Solarbio). The samples were separated by electrophoresis using 10% SDS-PAGE, and the protein samples were transferred to PVDF membranes using the Bio Rad semi-dry transfer system, blocked with 5% skimmed milk for 1 h, and the primary antibody was incubated at 4°C overnight. On the second day, the membrane was washed three times with TBST for 10 min each time, and then the HRP-conjugated secondary antibody was incubated at room temperature for 1 h. The membrane was washed three times with TBST and then blotted. The analysis and statistics of the results were performed using ImageJ software (NIH), and the histograms of the statistical results were produced using Graphpad.

### Immunohistochemistry

For immunohistochemistry, cryostat sections (14 μm) of mouse brain prepared as previously described were washed in PBS, and pre-incubated in PBS containing 5% donkey serum and 0.2% Triton X-100 for 30 min. Sections were then processed for immunostaining by overnight incubation at 4°C with primary antibodies including 6E10 (1:1000), GFAP (1:500), Iba1 (1:500), Vimentin (1:500), RTN3 (1:500), washed in PBS, and incubated in fluorophore-conjugated secondary antibodies (Invitrogen) diluted 1:1000 in PBS. Sections were washed, mounted in Fluoromount, and examined with a Zeiss LSM900 confocal microscope.

### Protein fractionation and Aβ ELISA

For extraction of Aβ fragments, we followed a fractionation protocol previously described. Dissected frozen mouse hippocampal tissue (9-month-old) was thawed on ice, and homogenized using a manual Dounce homogenizer, in Tris-buffered saline (25 mM Tris, 140 mM NaCl, pH 7.2– 7.6) containing protease and phosphatase inhibitor (MCE). Homogenates were then ultracentrifuged at 100,000 g for 1h at 4°C. The supernatant, which constitutes the soluble materials, was collected as the Tris-buffered saline fraction. The pellet of the Tris-buffered saline fraction was resuspended in RIPA buffer (Sigma-Aldrich) containing protease and phosphatase inhibitors by trituration 15 times with a 1ml pipette. Samples were incubated at 4°C for 1h with agitation and ultracentrifuged at 100,000 g for another hour at 4°C and were collected as RIPA fraction. The Tris-buffered saline and RIPA fractions were stored at −80°C until further analysis. The content of Aβ1-42 in the Tris-buffered saline and RIPA fractions was assessed using the Human Aβ1-42 enzyme-linked immunosorbent assay (ELISA) Kit (R&D) according to the manufacturer’s instructions. The concentration of Aβ1-42 was determined by the absorbance value detected using a microplate reader (BioTek Cytation 5, US) at 450 and 590 nm and then normalized to total protein concentration.

### Image analysis

For each mouse, at least five brain coronal sections were quantified. In each brain section, mosaic images in hippocampus were captured. ImageJ software was used to quantify positive signal area for the different markers. For Aβ, GFAP, Iba1 and Vimentin, percentage of positive signal area was normalized to total area of hippocampus. For RTN3, positive signal area within Aβ plaques was for quantification and expressed as percentage of total Aβ plaque area.

### Novel object recognition

The novel object recognition (NOR) test for mice consists of exposure training, followed by a short-term memory (STM) test 3 h after training, a long-term memory (LTM) test 24 h later. The objects are chosen based on similarities in dimensions and complexity. Tests are carried out in an acrylic box (20.32 × 40.5 × 16 cm LxWxH) that is sanitized, together with the objects, with 75% ethanol between each experiment. The time spent with an object includes direct visual orientation toward an object within half the body length of the object, sniffing, touching, or climbing on the object. The tests are video-recorded, and preference scores are calculated as time spent with the novel object minus time spent with the familiar object divided by the total time spent with both objects. Positive scores indicate preferences for the novel object; negative scores show preferences for the familiar object.

### Barnes maze

Twenty 5 cm diameter holes were evenly distributed around the edge of a 90 cm diameter circular platform. Three spatial cues of different shapes and colors were placed around the disc and kept constant during the experiment, and the platform was about 1 m above the ground with a strong light above it. Before the first training session, the mice were placed in an escape box for 1 min, and then the mice were placed in the center of the platform with a black box, which was removed after 10 s and recording began. During each training session, mice were free to explore the maze until they entered the escape box or for a period of 3 min. For each mouse, the escape box was always located below the same hole. Prior to each day’s experiment, the platform would be rotated 90° to avoid the influence of olfactory cues on the experiment. Mice were trained once a day for 5 days. A memory test was performed 3 h after the last training session. During the memory test phase, the escape box was removed and the mice explored freely in the maze for 90 s. The memory level of the mice was quantitatively assessed by calculating the exploration time spent in the target quadrant.

### Statistical analysis

Statistical Analysis Software (SAS Institute, Inc.) and GraphPad Prism (versions 4 or 8; GraphPad Software Inc, San Diego, CA, USA) were used for statistical analyses. Experimental data was collected from multiple experiments and reported as the treatment mean ± SEM. Statistical significance was calculated using the statistical analysis indicated in the Fig. legends. P value of less than 0.05 was considered statistically significant. *p<0.05, **p<0.01, ***p<0.001, ****p<0.0001 versus Control groups.

## Acknowledgements

The authors would like to thank Lei Wang, Jocelyn Jia, Yankui Fu and Shuo Zhang for technical and admin assistance. This work was supported by research grants to C.F.I. from Peking University, Chinese Institute for Brain Research, Beijing, and Swedish Research Council (Vetenskapsrådet, contract nr. 2024-03222); and a startup grant to M.X. from Swedish Research Council (Vetenskapsrådet, contract nr. 2021-01805).

## Author contributions

X.L. performed all experimental work, analyzed data and prepared a draft of the manuscript and figures; M.X. co-directed the project and corrected the manuscript; C.F.I. conceived the project, directed the research and wrote the final version of the manuscript.

